# Neuronal Decoding of Decisions in Multidimensional Feature Space Using a Gated Recurrent Variational Autoencoder

**DOI:** 10.1101/2025.08.20.671126

**Authors:** Charles Grimes Gerrity, Robert Louis Treuting, Richard Alan Peters, Thilo Womelsdorf

## Abstract

Recent advances in neuroscience enable recording neuronal signals across hundreds of channels while subjects perform complex tasks involving multiple stimulus dimensions. In this study, we developed a novel encode-decode-classify framework employing a gated recurrent variational autoencoder (VAE) to decode decision-making processes from over 300 simultaneously recorded neuronal channels in the prefrontal cortex and basal ganglia of monkeys performing a multidimensional feature-learning task. Using hierarchical stratified sampling and balanced accuracy, we trained and evaluated the model’s ability to predict behavioral choices based on neuronal population dynamics. The results revealed distinct neural coding roles, with anterior cingulate cortex (ACC) channels encoding decision variables collectively and prefrontal cortex (PFC) channels contributing individually to decoding accuracy. This approach demonstrated decoding accuracy for decisions in multi-dimensional feature space that is comparable to single-label decoding accuracy for lower dimensional problems, highlighting the potential of machine learning frameworks to capture complex spatiotemporal neuronal interactions involved in multidimensional cognitive behaviors. The code has been released in https://github.com/cgerrity/Neural-Data-Reading

## I. Introduction

A brain-computer interface (BCI) is a system that establishes a bidirectional link between an animal subject’s brain and a computer [1]. By bypassing the sensors and effectors a subject normally uses to interact with the world, a BCI allows for direct communication with the brain [2]. This bidirectional communication can range from reading out the activity of a single neuron and displaying the response of the subject to more complex systems where the BCI processes neuronal signals and modulates the subject’s behavior by stimulating the brain with a different signal. BCIs have been developed for many different domains, such as motor [3]–[8], speech [9]– [11], sensory [12]–[14], sensorimotor [15], and cognitive [16]– [22]. The most well known BCI would be the cochlear implant, allowing patients with auditory impairment to hear [23].

This work employs a value-based feature learning task. Value-based decision making involves comparison of expected value outcomes of potential choice options and choosing the option with the highest subjective value [24]. This can be viewed as choosing an option that maximizes utility. This type of learning requires encoding and updating of the value of relevant information based on feedback [25]. Decisions are made from this feedback, selecting the most probable rewarding outcome. Among the brain areas that are relevant for decision making and feature learning are the head of the caudate nucleus (striatum), the anterior cingulate cortex (ACC), and the prefrontal cortex (PFC) [17], [26]–[31]. These areas guide decision making by influencing information sampling, tracking subjective value, tracking reward prediction errors, and more.

The goal of this work was to develop a cognitive BCI that decodes decisions from neural activity recorded during a feature-learning task. Specifically, we present a decoding framework based on a variational autoencoder (VAE) with gated recurrent unit (GRU) to generate a low-dimensional latent representation of the neural dynamics. A long short-term memory (LSTM) network attached to the latent space units was trained to classify the feature dimensions encoded in the multiunit activity (MUA). This approach enabled us to analyze how neural signals across multiple brain areas relate to decision outcomes during a complex, real-world-inspired task.

## II. METHODS

### A. Behavioral Task

The experiment involved behavioral and neuronal assessment using a macaque monkey. All animal and experimental procedures were in accordance with the National Institutes of Health Guide for the Care and Use of Laboratory Animals, the Society for Neuroscience Guidelines and Policies, and approved by the Vanderbilt University Institutional Animal Care and Use Committee (M1700198-01).

Monkeys performed a feature learning task while probes recorded neural activity. For the task the monkey must learn a rewarded feature by choosing one of the three objects displayed on a computer screen [32]. To start a trial a monkey fixates on a black circle in the center of the screen. The trial starts and three different objects are presented. To select an object the monkey must fixate on it for 700 ms as shown in Figure 1A.

**Fig. 1.**
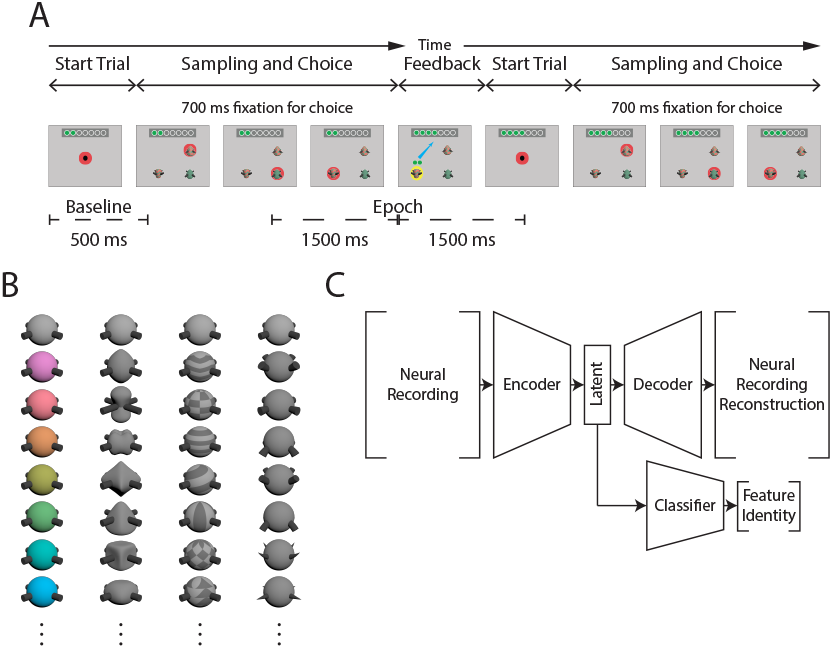
A) A token bar is shown to the monkey at the top of the screen indicating the number of tokens the monkey has received and how many until the bar is full. The monkey must fixate on the center dot to start a trial. The quaddles appear and the monkey chooses an option by fixating on it for 700 ms. If the correct quaddle was chosen a yellow halo appears around the object along with a visual indication of the accumulation of tokens. For incorrect trials a blue halo appears around the chosen quaddle followed by visual indication of the loss of accumulated tokens. If the number of tokens exceeds a set value, the monkey is rewarded with water and the tokens are reset to 0. B) The quaddles can have multiple dimensions depending on the number of non-neutral features it has. Each column is the four different feature dimensions, color, shape, pattern, and arms. The first row of each column is the neutral feature. C) This figure shows the network architecture used. The neural recording is input to the network-encoder to generate the latent space. The latent space is the input to the network-decoder and network-classifier. The network-decoder outputs a reconstruction of the neural recording. The network-classifier gives feature probabilities for each feature dimension.

One trial comprised exploring the objects until one object is chosen, triggering visual and auditory feedback indicating that the chosen object contained the target feature (resulting in reward) or not (unrewarded). The rewarded target feature stayed constant over a block of 25-50 trials. Neuronal recordings were performed during sessions in which monkeys performed 28-36 blocks.

Objects were so-called quaddles [33], [34] and examples are shown in Figure 1B. Each object has multiple feature dimensions each with a single feature value. These dimensions and features are aspects of the quaddle such as color (red, green, blue, etc.), shape (cylinder, pyramid, etc.), pattern (checked, lines, etc.) or arms (flared, pointed up, etc.). There are default null features that have been trained to be neutral (grey, sphere, no pattern, and arms straight out). The three quaddles that were presented in each trial, differed in the number of non-neutral features, either one, two, or three feature dimensions. This dimensionality was constant within a block of trials. Of all these varying features values, only a single feature was the target feature that was rewarded in each block, and the monkey had to learn this feature by trial-and-error.

For each choice the monkey received reward in the form of visual tokens. A token bar was shown to the monkey at the top of the screen indicating the number of tokens the monkey had received and how many until the bar was full. Monkeys earned either two or three token when they correctly chose the target object, and they lost one or three tokens when they chose an object without that target feature. The number of tokens earned for correct choices and removed for incorrect choices stayed fixed during a block, but varied between blocks. When the monkey accumulated eight tokens he received fluid reward and the token bar was reset to have 3 baseline tokens. Each block started with 3 tokens. After choosing an object there was a 500 ms feedback period during which the token bar was updated depending on the choice outcome, correct or error. A more detailed explanation of the task can be found in [32].

### B. Neural Recording and Preprocessing

MUA was recorded from six 64 channel linear silicon probes (Neuronexus). Details of the recording approach are described elsewhere [31] and involved placing recording probes prior to the start of a recording session into the ACC, PFC and the striatum using a software-controlled electrode drive system. Wide-band neuronal recordings were accomplished with an Intan Recording system, notch filtered at 60, 120, and 180 Hz to remove line noise and its harmonics. Subsequent preprocessing was done using the fieldtrip MATLAB library [35], and custom MATLAB code.

First, we removed compromised recording channels using principal components analysis (PCA) and K means clustering across trials. The number of means started at 2 possible groups and ended at half the number of channels on the probe (32). The distance measure between data points was squared Euclidean distance. If a channel was clustered by itself or a known artifact channel in more than half of the clusterings it was considered a bad channel.

Second, we re-referenced the signal by subtracting the median activity of the wideband data at each time point. Each probe-trial combination had a different median activity signal. This signal was then subtracted from each signal to remove the noise common across all the non-artifact channels. This method of re-referencing was found to be better for spiking data instead of average re-referencing which removes the average activity of each probe [36], because single spikes would be averaged into a smaller amplitude spike and then appear on each channel.

Third, to obtain continuous MUA the re-referenced trials were band-pass filtered (0.75-5kHz), rectified, low pass filtered (0.3kHz), resampled at 1 kHz and smoothed with a 200 ms gaussian window. This generates a measure of MUA that maintains small amplitude spikes [37].

Fourth, the neuronal data were segmented into individual trials. Each trial was aligned to the time of the object choice and then 1500 ms before and 1500ms after were taken as the whole epoch, as shown in Figure 1 A. Additionally, a baseline epoch was identified as the first 500 ms of the trial, which contained the start of the trial period and was used as a baseline for the activity.

Fifth, we detrended the data, using regression of the average activity of the baseline period preceding the choice of an object in each trial. The regression trend across trials was removed from both the baseline and decision epoch.

Sixth, we z-normalized the data of each MUA channel by calculating the mean and standard deviation in the baseline period across the entire session per individual channel and subtracted each trial by that mean and dividing it by the standard deviation.

Seventh, we used regression analysis to identify MUA channels that were not modulated during any epoch of the trial and removed them from further processing [38]. This analysis determined the encoding of the task variables in the activity. Each trial of the task had a different set of task variables associated with it. These were used as the independent variables in the regression as shown in Table I. The dependent variable was the continuous MUA. For each time point there were several observations, represented by each trial. The regression indicated whether the MUA at each time point was modulated by the task. We removed channels when there were no 50 ms long segments containing regression coefficient exceeding a significance threshold of 0.05 was used for identifying a significant regression model.

**TABLE I.** VARIABLES USED FOR REGRESSION ANALYSIS

| Variable | Type | Description |
| --- | --- | --- |
| $y$ | Scalar | Continuous MUA |
| $x_{reward}$ | Binary | Rewarded vs Unrewarded |
| $x_{learned}$ | Binary | Learned vs Unlearned |
| $x_{attention}$ | Two Binary | Dimensionality |
| $x_{gain}$ | Binary | High vs Low Gain |
| $x_{loss}$ | Binary | High vs Low Loss |
| $x_{previous}$ | Binary | Previously Rewarded |
| $\beta_0$ | Scalar | Intercept |

Eight, after these processing steps the individual areas from each trial were concatenated along a new dimension, brain area recording location. The total number of recording locations across areas, A, was the maximum number of different probes recorded. The maximum number was 6 probes, two in each of the areas, ACC, PFC, and striatum.

Let **M**_*i*_ *∈* ℝ^*C×T*^ for i = 1, 2, …, A be a set of A matrices, each with dimensions C by T. C is the number of channels. T is the number of time points. The final trial output is *X* where *X ∈* ℝ^*C×T×A*^ is a tensor with elements: *X*_*c,t,a*_ = (**M**_*a*_)_*c,t*_ *∀ c ∈ {*1, 2, …, *C }, t ∈ {*1, 2, …, *T }, a ∈{* 1, 2, …, *A }*. In other words, the resulting tensor X is constructed by stacking the A matrices **M**_*i*_ along a new third dimension, where each matrix becomes a slice of the tensor.

The order of the areas is also identical across all the *X* where the first two matrices represent recordings from the ACC, the next two from the PFC, and the final two from the striatum. For sessions where there was not a recording from an area, a matrix of empty values was added in its place. This maintained the final data with the same shape *X ∈* ℝ^*C×T×A*^, but did not introduce uncollected data.

### C. Hierarchical Stratified Sampling

To assess the decoders ten-fold cross validation was used [39], [40]. Each fold represented an equal portion of the data. The folds were used to construct training, validation, and testing sets as 80%/10%/10% splits. Repeated cross validation was not performed as it would add too much to the computational cost as well as leading to a bias in the estimate of model performance [39].

The target distributions are imbalanced, meaning there is an over representation of classes. The majority of chosen features are the neutral feature because every object has at least one neutral feature. This leads to a class imbalance across the four labels.

Methods address class imbalance in training by over-sampling the minority classes, under sampling the majority class [41]–[43], by modifying the loss function to penalize minority classes more to balance their infrequency [43], or by stratified sampling [44]. Here, we addressed class imbalance through stratified sampling, which has been shown to improve models by ensuring that each fold has the same distribution of features as the original dataset. We partitioned the data into subsets called strata [40], [45], which maintained the same proportion of these strata [45], [46]. Strata for single label classification are typically the target labels, whereas for multi-label classification the strata could be the combinations of labels also known as labelsets [46].

Using stratified sampling classification accuracy on *k*-fold training has improved bias and variance estimates of accuracy compared to random partitioning of the data [40], [46]. Additionally, stratified sampling can reduce the number of partitions that do not have any examples in the possible labelsets [46]. The strata should have an equal representation of the four feature dimensions. One stratum with an over representation of a class could skew the trained model [44]. Not only is it important to keep the distribution of the chosen features consistent, but it is also important to keep the distribution of the trial variables consistent, which included the motivation condition (high versus low number of tokens gains/lost), the outcome of the trial (correct vs error), of whether the trial occurred during learning or when learning completed. While the trial variables are not the target of the models, they still play a crucial role in what information is found in the neural data. Trials before the subject has learned the rewarded feature will be different than after given the importance of the areas for learning. This may also be true for the gain and loss conditions because they affect the motivational state of the subject [47].

To produce the strata the data is split into ten folds using custom stratifying code. Named here as hierarchical stratified sampling. This method assigns a group label to each trial. Then partitions are generated by randomly sampling from each group, ensuring that trials in the same groups end up in different partitions. For this method the variables for splitting the trials are ranked. The rank list was as followed:

1. Shape, pattern, color, and arm type
2. Recording probe brain area
3. Outcome
4. Gain and loss condition
5. Learning status
6. Session name

The method starts by identifying all the unique combinations of the top most rank. This would mean all the feature combinations that appear in the data. Then for each group, if it has more elements than the number of folds (10) then it will move to the next item on the rank list. This group will be split by the unique combinations of this rank. Then again, if each combination has more elements than the number of folds the group moves to the next rank down the list, otherwise the group is given a unique label. This continues until the whole rank list has been traversed or there are not enough elements in the groups to continue moving down the list. This gives each trial a group number which is used to randomly assign each element of a group to a different partition.

### D. Metric

In addition to addressing class imbalance in training with hierarchical stratification, we addressed class imbalance in the model evaluation by using balanced accuracy on the BCI decoder output.

Accuracy is misleading when looking at an imbalanced dataset [43], [48]–[51], because a most common dummy classifier will have an accuracy equal to the majority class frequency. If balanced accuracy is used then no matter the class imbalance the measure is unchanged. The issue of using accuracy could be mitigated by listing the chance measure as well [11], [21]. However, the selection of how chance is calculated may differ depending on the specific use case. To generate a chance estimate a dummy classifier is selected, which is a classifier that attempts to classify the data naively [52]–[54]. Two types that would be relevant here are the most frequent and stratified. The most frequent dummy classifier will always output the most frequent class. If there are two classes and *p*_*majority*_ represents the proportion of the majority class and A represents the accuracy then A = (*p*_*majority*_)(1) + (1 *− p*_*majority*_)(0) = *p*_*majority*_. This will lead to A = |*p*_1_ *−*0.5 |+ 0.5. The stratified classifier will output random classes with the same frequency as the training set. If there are two classes and *p*_1_ represents the proportion of class one 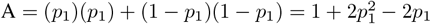 Both dummy classifiers will give different estimates of chance level for accuracy, leading to different improvements described by using a classifier depending on the chosen dummy classifier.

Balanced accuracy for binary cases is the average of sensitivity and specificity [43], [48]–[51]. For multi-class classification this becomes the accuracy for each class, also described as the class recall.

Balanced accuracy has two main features: it maintains a stable estimate of chance regardless of class imbalance and it equalizes the choice of dummy classifier. For the two class case, the balanced accuracy from the most frequent dummy classifier is BA = (0.5)(1) + (0.5)(0) = 0.5 and the balanced accuracy from the stratified dummy classifier is BA = (0.5)(*p*_1_) + (0.5)(1*− p*_1_) = 0.5

Balanced accuracy addressed the issues that arise in imbalanced datasets, but another issue is the identification of what is a good value of the metric. To this end metrics are compared to chance levels [11], [21], which helps distinguishing a poor classifier from a good classifier. Specifically, the metric can be chance corrected by scaling the value so that 0 represents chance level accuracy and 1 represents perfect classification [55]–[60]. Balanced accuracy has a stable chance level allowing for chance correction to be effective [61].

We calculated scaled balanced accuracy, 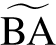, as the average class recall of a decoder. Let *M* be the number of feature dimensions and *N*_*j*_ be the number of feature values in dimension *j*. Given a decoder, *d*, over the set of features per dimension, Equation 1, the true positives (*TP*) and false negatives (*FN*) are calculated for each feature, *F*_*i,j*_, with balanced accuracy reflecting the average of the class recall, as shown in Equation 2. To extend balanced accuracy to multiple labels, the balanced accuracy for each dimension, *D*_*j*_ is averaged, Equation 3.

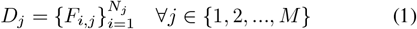

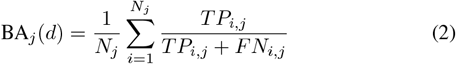

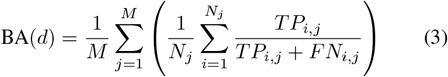

There is an issue of what is a good value. To this end metrics can be compared to chance levels [11], [21]. This helps distinguishing a poor classifier from a good classifier. Specifically, the metric can be chance corrected by scaling the value so that 0 represents chance level accuracy and 1 represents perfect classification [55]–[60]. Without using a chance corrected metric it is difficult to compare results on different sets of data [60]. Balanced accuracy has a stable chance level allowing for chance correction to be effective [61]. Here the balanced accuracy is scaled to a chance decoder, *c*, Equation 4.

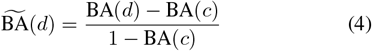

To address the issues from imbalanced data, choice of dummy classifier, ease of understanding, and comparison to other work, balanced accuracy was selected and scaled to chance. This measure is used for all reports of accuracy in this paper. The neural data is a time series so where possible the time series of accuracy is presented and when single values are presented peak accuracy at any time is used.

## E. Decoder Architecture

Previous work in BCIs has shown that encoder-decoder networks perform well [6], [8], [62]–[69]. This type of network is able to remain stable over multiple chronically recorded sessions [67], which will help to apply this work to multiple acutely recorded sessions.

Autoencoders are a type of encoder-decoder network where the output attempts to match the input [70]. They are able to compress data or reduce dimensionality by progressively smaller and smaller layers in the encoder down to a latent space followed by a progressively growing decoder to reconstruct the input from the latent space. They are able to be trained in a self-supervised manner [8], [71]. An autoencoder was selected to provide a non-linear compression of the data into a latent space. VAEs are a type of autoencoder, where the latent space is represented by a probability distribution with learned parameters [72]. Using a VAE, points in latent space can smoothly transition from one representation to another without abrupt changes.

The overall architecture used for the artificial neural network (ANN) based decoder can be seen in Figure 1 C. The core architecture is similar to previous work [6], [67], [68], but was adjusted to accommodate the multidimensional feature input.

The network has three parts, encoder, decoder, and classifier. The encoder has sequential layers with half the number of units to reduce the data down. The last layer has twice the number of units to account for the mean and variance components. At the end of the encoder is a bottleneck where the last layer is repeated for the same number of units according to a hyper parameter controlling the depth of the bottleneck. The decoder contains layer of increasing size with an initial sampling layer to generate a variational autoencoder. The classifier has input size equal to the output size of the encoder. There is a separate classifier network for each feature dimension. Each classifier can classify the neutral feature of the dimension or a non-neutral feature. Hyper parameters for the model were chosen using a sweep over parameter space.

The input to the network was the windowed MUA trial data. Each trial of the data *X* where *X ∈* ℝ^*C×T×A*^, was segmented by trial time according to the data width parameter. This window slid over the data with a specified stride. This generated a new input *X*[*s*] where *X*[*s*] *∈* ℝ^*C×W×A*^. Here *W* represents the size of the window and there are *S* windows, which is the maximum number of windows using the specified stride. Comparing this to video data, the *C* and *W* dimensions would be analogous to the spatial dimensions. The *A* dimension would be the channel dimension. Lastly the *S* would be the time of each frame.

### F. Alternative BCI Decoding Architectures

Other model types were tested: feedforward, GRU, LSTM, convolutional, residual network (ResNet), and multi-filter convolutional. Some models of BCI decoders have been feedforward networks for movement [6], [67]; GRU and LSTM based networks for speech [10], [62], [73], [74]; and convolutional networks for functional magnetic resonance imaging (fMRI) visual decoding [71]. For each of the models corresponding blocks as shown in Figure 2 were repeated. Common to each block were dropout and normalization layers to improve generalizability and training stability.

**Fig. 2.**
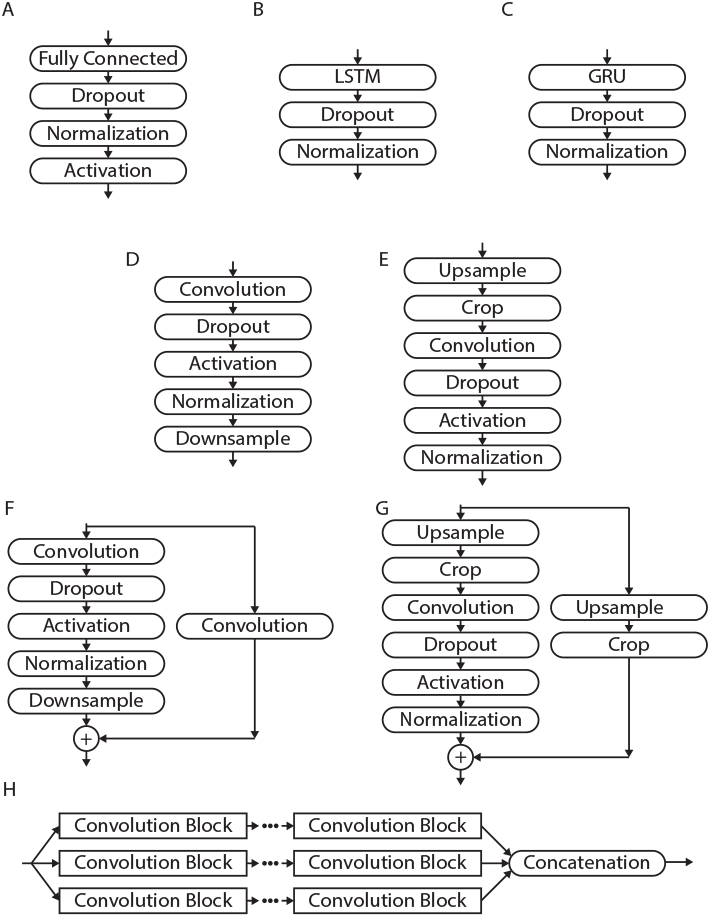
This figure shows the blocks used to for the different network models. A) shows the layers for each feedforwad block. B) shows the layers for each LSTM block. C) shows the layers for each GRU block. shows the layers for the convolutional blocks for the encoder and shows the layers for the decoder version. F) shows the layers for the ResNet encoder blocks and G) shows the layers for the ResNet decoder blocks. H) shows the structure for the multi-filter model using convolutional blocks with different filter sizes.

The feedforward blocks, Figure 2A, began with a fully connected layer and was followed by a dropout layer, normalization layer, and activation layer. The activation for the feedforward blocks were rectified linear unit (ReLU). The LSTM and GRU blocks, Figure 2B and C, began with the specified recurrent layer and was followed by a dropout layer and normalization layer.

The convolutional blocks started with two-dimensional convolution with filter size equal to 30% of the channels and window size (*C × W*). After convolution, the next layers were dropout, leaky ReLU, and normalization. Convolutional blocks reduce their output size by down sampling instead of reducing the number of units. This was performed by adding a 2*x*2 max pooling layer with stride of 2. This takes the maximum value in a 2*x*2 grid as the output, reducing the size by half. For the network-decoder version of the convolutional block, the alternative was upsampling. This was performed at the start of the block using transpose convolution with filter size the same as the network-encoder. This identifies the array that when convolved with the filter generates the input [75]. Cropping was applied to match the corresponding encoder size at the given layer then two-dimensional convolution was applied for the desired number of filters.

The ResNet block was identical to the convolutional blocks, but with an added skip connection. The skip connection can help with deep networks and the vanishing gradient problem, where the gradient is so small that it does not meaningfully change the weights [76], [77]. A separate path was constructed around the convolutional block that used either convolution or transposed convolution with stride 2 for the network-encoder or network-decoder respectively. This was added to the original convolutional path. Without the application of the activation the network can learn the skip path as the identity and the non-skip path as a residual that adds to the identity.

Lastly the multi-filter block was a set of convolutional blocks with different filter size percentages, 20%, 30%, and 40%. These processed identically to the convolutional version, but there was a final step where the outputs from each filter were concatenated.

### G. Training

Each model was trained using the MATLAB deep learning toolbox [78] and custom MATLAB code. Training leveraged the resources provided by Vanderbilt Advanced Computing Center for Research and Education (ACCRE), a collaboratory operated by and for Vanderbilt faculty.

At the beginning of each training session, the data was partitioned into training, validation, and testing sets using the hierarchical stratification procedure described in section II-C. For all partitions the data was normalized. First each channel was z-scored according to the mean and standard deviation across all trials. This generated data where each channel was centered on 0 with unit standard deviation. Next the data was min-max scaled to lie in the interval [*−*1, 1]. Lastly, because the min-max scaling changed the mean of the data away from 0 the data was recentered so the mean across all of the data was 0 and it was also scaled so that the standard deviation across all the data was 0.25. This generated data that had most data points in the range of [− 0.5, 0.5]. This was done to allow for sufficient range of the data in case the min-max scaling compressed the majority of the data into a very small range.

All channels that were removed during the preprocessing and processing steps were set to 0 for all time points. These channels were not included in the normalization procedure, so they did not affect the calculations of mean, standard deviation, minimum, and maximum. Next for the training set only, data augmentation was applied to generate new samples with similar characteristics to promote generalizability. Random noise was added to each channel each time the data was used in training. The noise had three components, channel offset, random walk, and white noise [74], [79]. The channel offset was a constant offset added to a given channel. The white noise was Gaussian noise. The random walk was the cumulative sum of a separate white noise generator. After generating noise for each channel, it was smoothed with a 50 ms Gaussian window to obtain noise with similar frequency characteristics to the data. Without this final smoothing the noise added was entirely at a different frequency to the data. This would lead the model to identify and remove high frequency noise, instead of seeing each training sample as new examples. Because high frequency noise was not present in the unaltered data, this would add an unnecessary parameter for the model to learn that would not contribute to generalizability. The standard deviations for each type of noise were 0.3, 0.15, and 0.007 for channel offset, white noise, and random walk respectively.

For training the networks themselves, the first step was getting the class weights, *w*_*i,j*_. This was taken as the inverse frequency of each class, *j*, for each dimension *i*. This prevents the model from emphasizing the most frequent feature, which is the neutral feature.

The learning rate for each epoch was determined by a ramping learning rate followed by learning rate decay. The ramping period was set to 5 epochs. During this period the learning rate was linearly increased from 0 to the initial learning rate. This was implemented to prevent large changes in weights that was occurring when the learning rate started at the initial value. The learning rate decay occurred every 30 epochs and decreased the learning rate by 90%. This allowed for faster learning at initial epochs followed by slower learning when approaching the final model. For each epoch all data in the training set was used by splitting it into mini-batches. The mini-batches were randomly partitioned from the training set at the start of each epoch.

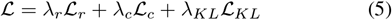

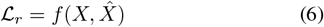

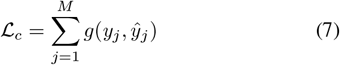

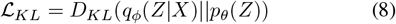

For each iteration the model loss was calculated using Equation 5, where *ℒ* is the total training loss which is a weighted sum of reconstruction loss *ℒ*_*r*_, classification loss *ℒ*_*c*_, and Kullback-Leibler divergence *ℒ*_*KL*_. This loss has a reconstruction component, Equation 6, which is used to measure the difference between the input, *X*, and its reconstruction, 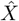. The function for reconstruction loss was mean squared error, Equation 9, averaged over the mini-batch. For each channel in the continuous MUA that was removed and set to 0, the contribution to loss from that channel is set to zero. This prevents the network from trying to fit the reconstruction to these removed channels. The next component is the classification loss, Equation 7. This measures the difference between the one-hot encoded true label, *y*_*i,j*_, and the predicted label, *ŷ*_*i,j*_ for the *j*-th feature out of *N*_*i*_ features for the *i*-th dimension. Cross-entropy was chosen as the classification loss function, Equation 10, averaged over the mini-batch. The contribution of each class to the loss is weighted by *w*_*i,j*_. The last term is the KL loss which comes from the variational component, Equation 8. This loss measures the difference between the latent distribution, Equation 11, described by the encoder output, and a standard normal using the KL divergence, Equation 12. Here, *L*, represents the size of the latent space. *µ*_*e*_ and 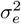 represent the outputs from the encoder.

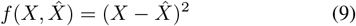

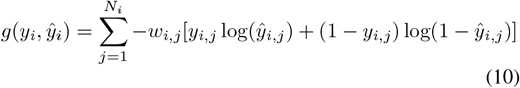

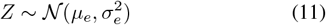

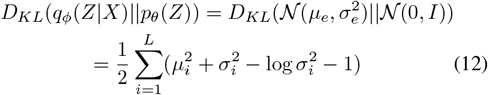

For each loss component there was a weighting *λ*. This was used to parameterize the contribution of each of the loss components [6], [65], [67], [80]–[82]. Here the weighting was scaled according to the current loss similar to other BCIs [6], [65], [67]. Equation 13 shows that each weighting was scaled by a weight factor, *ω*_*i*_, and the loss at the first iteration of each epoch similar to previous studies [6]. The ratio loss between each component was roughly maintained at the ratio between their weight factors and resets at the start of each epoch.

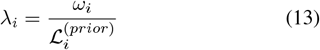

The gradient for each network component was calculated using backpropagation through time. Certain models and parameter combinations generated a large memory footprint. This led to training slowdowns or failed runs. This was ameliorated by using gradient accumulation. This processed smaller partitions of the mini-batch to obtain loss and gradient values, and then combined them. Processing in parts allowed for the more accurate gradient estimation obtained by larger mini-batch sizes without the problems of a large memory footprint. Another gradient issue was the exploding gradient. To prevent this a threshold was applied to the gradient for each component. The *L*_2_ norm of the gradient was used to scale the gradient to the maximum set by the gradient threshold parameter, but only if it surpassed the parameter. By setting an upper limit of the gradient threshold parameter, the gradient was forced to stay below that and not explode. This gradient clipping was set to 100. The gradient was used to update the model weights using the optimizer. Both stochastic gradient descent and ADAM were used as optimizers. In addition, L_2_ regularization was applied. This prevented weights from growing very large, preventing overfitting.

Scaled balanced accuracy was used to monitor the accuracy of the model. The output of the network-classifier was a probability distribution over the features of each dimension. The highest probability feature was selected as the predicted feature for each dimension. Other work has added language interpreters to the output of their models to generate sensible speech from phoneme probabilities [10], [74], [83]. Similar to this interpreter approach, a simpler interpreter was generated for the quaddles. Each quaddle could be at most 3-dimensional and at least 1-dimensional. The interpreter took the feature probabilities and generated a possible quaddle as described.

However, if an illegal quaddle was generated, then one of two changes were made. First the neutral confidence, the difference in probabilities between the neutral feature and each non-neutral feature was obtained. If too many neutral features were detected, then the feature with the lowest neutral confidence was changed from neutral to the corresponding non-neutral feature. If too many non-neutral features were detected then of the four non-neutral features, the one with the highest neutral confidence was changed to the neutral feature. The interpreter generated legal quaddle outputs that the subject could have actually chosen. The output object at each time point was used to calculate the balanced accuracy over time.

The last step of training was the calculation of the validation loss and accuracy. These values were used to determine the optimal network. For full network training the maximum validation accuracy at any time point was used to select the best performing model.

## III. Results

### A. Comparison to Alternative Approaches

Parameter selection was tested on different values for the various network and training parameters. For parameter selection each possible combination was trained for 100 epochs. This was chosen to give each set of parameters enough epochs to train, but not too many that the training time would be infeasible. Optimal network hyper parameters were determined by selecting the et of parameters that gave the highest validation accuracy. Figure 3 shows the chosen hyper parameters generated the optimal model given testing set results.

**Fig. 3.**
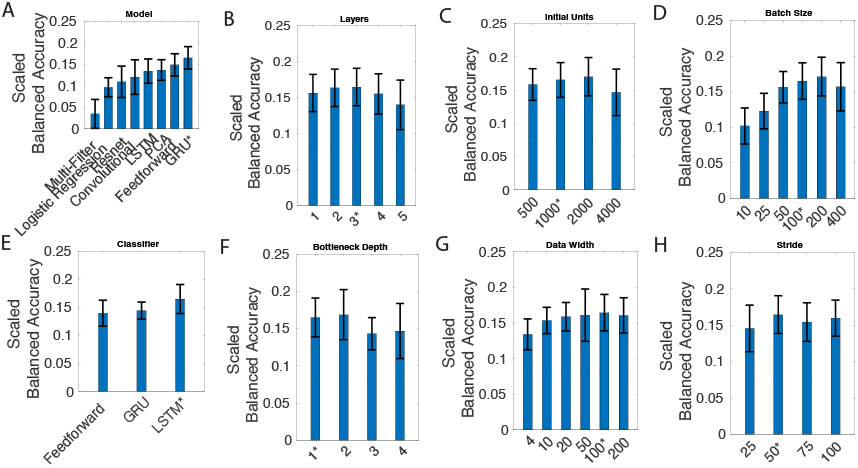
This figure shows the optimization of network parameters. A) represents the network-encoder and network-decoder blocks uses. B) refers to the number of layers for the network-encoder and network-decoder. C) refers to the size of the first layer. D) refers to the number of training example for each iteration. E) represents the network-classifier model used. E) is the depth of the bottleneck layers of the Encoder. G) refers to the size of the time dimension for each window of the input data. H) is the time shifted across the data for each window. indicates the parameter that was used in the optimal model.

Figure 3A shows that the chosen model remained among the best on an unseen testing set. The logistic regression uses only linear assumptions about how the neural data maps to chosen features and does not outperform the chosen model. Additionally, the PCA model, using linear assumptions about the reconstruction to generate the latent space, does not outperform the chosen model. Both of these results indicate that the neural data is best represented by a non-linear mapping to a latent space and a non-linear mapping for classification.

### B. Accuracy

The output of the optimal model is presented across the ten hierarchically stratified folds using the scaled balanced accuracy metric. In Figure 4 the peak testing accuracy is shown. This was taken as the average of the maximum accuracy at any window for each fold. In Figure 5 the average of the accuracy at each time point was calculated across folds, where all the windows are aligned. The accuracy was also calculated by splitting the testing set by the trial variables to determine which types of trials were best decoded

**Fig. 4.**
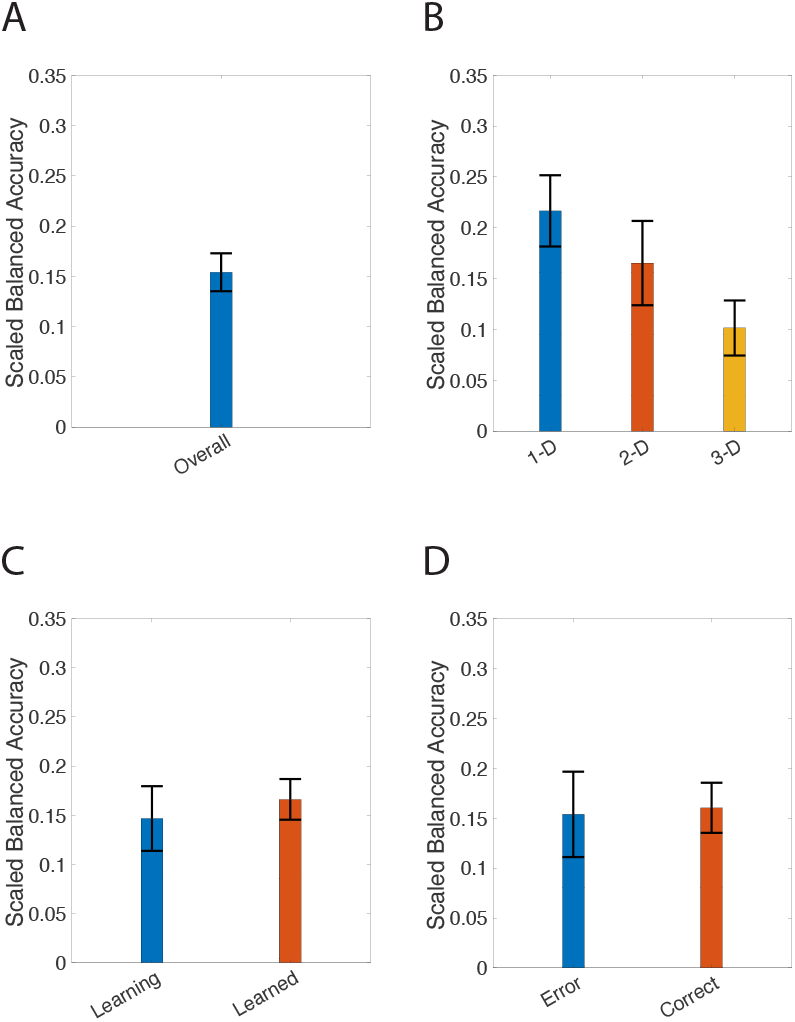
This figure shows the peak testing accuracy of the network across trial variables. A) The accuracy of the model on all testing trials. B) The accuracy for testing trials split by attentional load. C) The accuracy for learning status. D) The accuracy for different trial outcomes. Mean and 95% confidence intervals are presented.

**Fig. 5.**
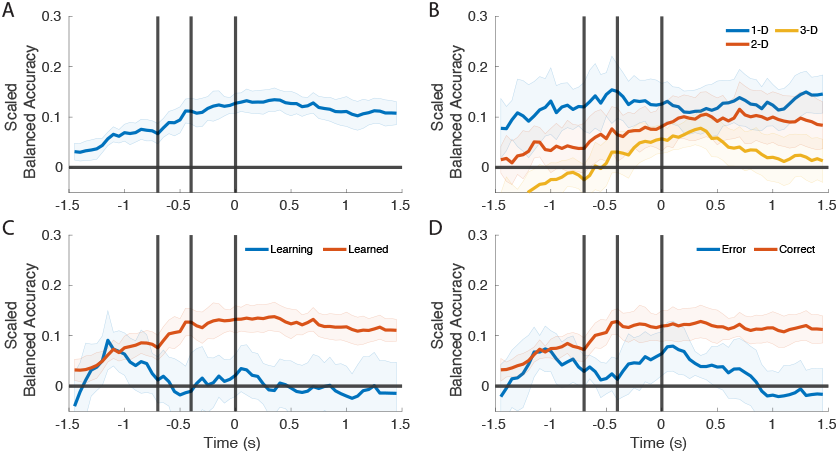
This figure shows the accuracy of the network across trial variables across time. A) The accuracy of the model on all testing trials. B) The accuracy for testing trials split by attentional load. C) The accuracy for learning status D) The accuracy for different trial outcomes. Mean and 95% confidence intervals are presented.

The peak overall accuracy was around 15%, Figure 4A. Figure 4B shows the decoder had a higher accuracy for decoding the trials in which features of only one dimension varied (1D) compared to trials with features of 2 or 3 dimension varying (2D, 3D). Figure 4C shows that peak accuracy on learned trials is similar to peak accuracy on learning trials. Whether a behavioral choice was a correct or erroneous had only a small effect on the peak accuracy (Figure 4D).

### C. Accuracy Over Time

Figure 5 examines the time course of accuracy over the trial. Across most of the conditions. Accuracy tends to increase across the trial remaining flat during the feedback period. Figure 5A shows the overall accuracy. There is an increase in accuracy starting before the subject has fixated on the object and continues increasing throughout the fixation period, suggesting that neuronal activity predicts the object will be fixated and chosen. Figure 5B shows the effect of dimensionality. The accuracy for 1-D trials remains fairly flat from start to end of the trial. The accuracy for 2-D trials starts to increase from the start of the trial and becomes significant at the start of fixation. The accuracy for 3-D trials similarly increases from the start of the trial, but is below chance until it becomes significantly above chance roughly 200 ms before the decision is recorded. The accuracy increases into the feedback period where it then drops off to chance. Figure 5C shows the effect of learning status on accuracy. Like overall accuracy, learned trials have above chance decoding throughout. For learning trials there is a peak in decoding accuracy before final fixation to choose the object. Lastly, Figure 5D shows the accuracy for trial outcome. The accuracy is highest for correct trials and appears similar to the overall accuracy. Error trials are above chance briefly at the start and briefly around the recording of the decision.

### D. Importance Analysis

We evaluated the contribution of each channel to decoding using an importance analysis that removed individual channels to judge their importance for the overall decoding accuracy [10], [73]. Importance analysis was performed by zeroing out a channel or groups of channels [4], [10], [73] and comparing the accuracy with all channels to accuracy of channels removed. We deployed three different approaches to evaluate how individual channels of the 321 possible channels contributed to the decoding accuracy, random, ranked, and sequential selection of removed channels. Random selection entailed testing 500 random combinations of removed channels at each number of removals and selecting the combination that produced the lowest peak accuracy. For ranked selection, the channels were ranked based of their decrease in peak accuracy from single channel removal. Using this rank list the removal was determined by selecting the highest-ranking channel, obtaining the accuracy, then removing the next highest ranking channel, and so on. For both removal methods the peak accuracy was chosen as the peak at any one time point averaged over the folds. Sequential selection starts by selecting the single channel with the largest drop in accuracy. Then the next set of most important channels is selected by including the channels from the previous list and testing all the remaining channels with them.

Figure 6A, B, and C shows the accuracy across folds for the different number of removals. Figure 6D, E, and F show the percent of removed channels per area as the number of channels removed increases relative to equal removal from each area,corresponding to the line connecting [0, 0] to [321, 1]. This shows the importance of probes from each area to the selection method. Probes from areas with high importance would increase earlier than areas that were selected later. Figure 6G shows the peak accuracy per method as a function of the number of removed channels. This would correspond to identifying the peak in Figure 6A, B, and C for each number of removals. Figure 6H is the time associated with the peak accuracy. Lastly, to get a measure of the importance of each area according to the given method the values from the percent relative to mean are averaged (Figure 6I). If an area is more important it will have a higher value.

**Fig. 6.**
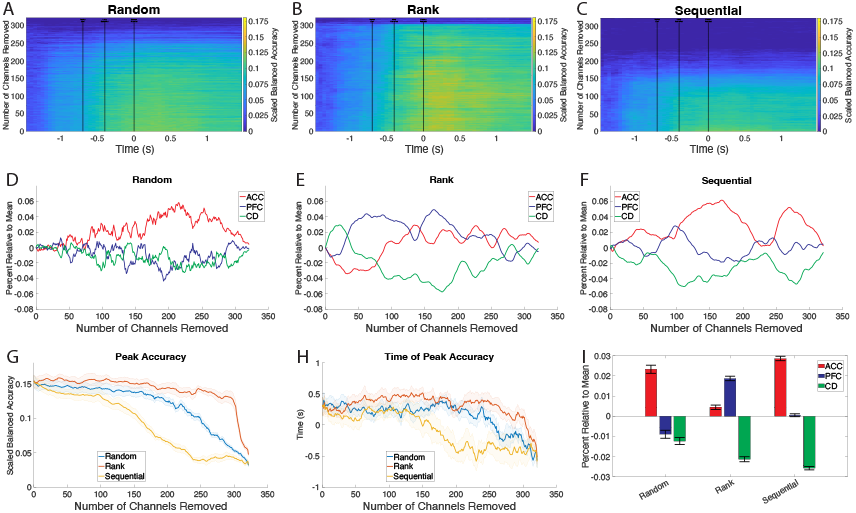
This figure shows the results of the importance analysis. A, B, and C) show the balanced accuracy over time for each number of channel removals. As more channels are removed the accuracy decreases. The removal order is determined by random and rank selection. D, E, and F) show the percentage of channels removed from each area compared to what that percentage would be if channels were removed equally across the three areas. G) shows peak accuracy over the number of channels removed for each method. As channels are removed accuracy decreases. H) shows the time of peak accuracy as channels are removed. I) shows the average relative percent of channels from each area. This shows which areas are most important for decoding.

### E. PCA Comparison

PCA was used to generate a latent space instead of using the encoder-decoder model. The latent size for this kind of model was similar to the chosen size of 500 for the network-encoder and network-decoder (MEAN = 471.8; SD = 0.4). Figure 3A shows the peak accuracy when compared to the various methods. The PCA based model is not able to capture the nonlinearities that are present in the representation of features.

## IV. Discussion

This work introduced a new cognitive BCI for multi-label multi-class decoding of decisions with above chance accuracy from high-dimensional neuronal recordings. The implementation of hierarchical stratified sampling addresses class imbalance systematically, ensuring robust model evaluation across diverse trial conditions. The effects of trial difficulty and outcome in the temporal dynamics of the accuracy reflected the expected behavior. Lastly the machine learning tool of feature importance translated the decoder structure to neural space.

Previous BCI work has focused primarily on either multi-label or multi-class decoding. Here the decoding problem involves both, multiple feature dimensions (labels) and multiple feature values per dimension (classes). Scaled balanced accuracy showed the utility of the decoder without requiring knowledge of chance or selection of a chance decoder. The chosen ANN decoder was able to achieve around 15% peak accuracy. This is similar to 20%, converting their work to scaled balanced accuracy, achieved in previous work [28] in a single label binary decision decoding task using the ACC. Although peak decoding accuracy is modest, it reflects a meaningful advance given the inherent complexity of the decoding challenge, multi-label multi-class classification on high dimensional data.

The type of architecture we implemented is similar to other architectures that generate a latent space for use in decoding from neural data [84]. The method does not try to isolate variables within the latent space but generates a latent representation that can best be used for further classification. A latent space that successfully isolated the various features of a decoded object would be analogous to an object decoder, isolating the combinations of features within the latent space. This would lead to sparsely represented objects and is not as feasible with features that do not have a continuous representation such as velocity or position.

The accuracy of the decoder reflected expectations of the task variables. For increased difficulty of the task the accuracy decreased, indicating that rising behavioral difficulty goes along with rising difficulty to accurately decode task variables.

Feature importance is used in many machine learning problems to identify relevant inputs to models. Here feature importance was used to quantify two different types of contributions of individual channels to the decoding performance. First, the overall joint contribution of a recording channel from a brain area was evaluated using random and sequential sub-selection of channels. We found that channels from the ACC had highest joint contributions. These represent joint contributions because the order of removals involves the contribution of the selected channel and the other channels that are considered with it. Second, the most informative channels for peak decoding accuracy were identified based on removing channels ranked according to their individual accuracy. We found that the probes with the highest individual contributions were from the PFC. Because this method looks at each channel in isolation it only considers how each channel contributes individually to the accuracy of the model. The discrepancy of the random, rank, and sequential selection of channel removal points to a difference of how channels jointly contribute to decoding performance in the ACC, while in the PFC, individual channels contributed to decoding without interacting with other channels. Because the sequential peak accuracy is lower than the rank or random peak accuracy, the importance of each channel is best characterized by joint contributions, instead of their individual contribution.

This work serves as support for further work in interpretable latent representations, temporal dynamics of cognitive processing, and brain region specific contributions to learning. Insights into these aspects of neuronal encoding and decoding of cognitive variables will be invaluable for the development of cognitive BCIs.

## Acknowledgment

We thank Adam Neumann for invaluable help with the experiment. This work was supported by the National Institute of Mental Health (R01MH123687). The funders had no role in study design, data collection and analysis, the decision to publish, or the preparation of this manuscript.

